# Blood gene expression-based prediction of lethality after respiratory infection by influenza A virus in mice

**DOI:** 10.1101/2020.10.27.357053

**Authors:** Pedro Milanez-Almeida, Andrew J. Martins, Parizad Torabi-Parizi, Luis M. Franco, John S. Tsang, Ronald N. Germain

## Abstract

Lethality after respiratory infection with influenza A virus (IAV) is associated with potent immune activation and lung tissue damage. In a well-controlled animal model of infection, we sought to determine if one could predict lethality using transcriptional information obtained from whole blood early after influenza virus exposure. We started with publicly available transcriptomic data from the lung, which is the primary site of the infection and pathology, to derive a multigene transcriptional signature of death reflective of innate inflammation associated with tissue damage. We refined this affected tissue signature with data from infected mouse and human blood to develop and validate a machine learning model that can robustly predict survival in mice after IAV challenge using data obtained from as little as 10 μl of blood from early time points post infection. Furthermore, in genetically identical, cohoused mice infected with the same viral bolus, the same model can predict the lethality of individual animals but, intriguingly, only within a specific time window that overlapped with the early effector phase of adaptive immunity. These findings raise the possibility of predicting disease outcome in respiratory virus infections with blood transcriptional data and pave the way for translating such approaches to humans.

## Introduction

Influenza A virus (IAV) infection of the respiratory tract can lead to severe immune activation and lung tissue damage in mice, ferrets, macaques and humans (*1–6*). Intrinsic virulence, replication capacity and initial infectious dose, together with the host’s genetic background, immune status and overall health, determine the extent to which host immunity is triggered (*7, 8*). Substantial evidence indicates that lethality is associated with an excessive innate immune response, with lung dysfunction arising from epithelial and endothelial damage induced by infiltrating leukocytes, in particular monocytes and neutrophils (*9–13*).

Our previous work in a murine model of infection with IAV uncovered clusters of co-regulated genes in the lung associated with lethal influenza infection; one of those lethal clusters was highly associated with an overwhelming neutrophil response and, consistently, early post-infection partial neutrophil depletion rescued animals from lethality, establishing a direct link between the innate response in the tissue and death (*13*). While that earlier work focused on gene expression signatures in the infected pulmonary tissue, here we sought to identify a blood-based signature for prediction of lethality.

Previous attempts to develop gene expression signatures in blood in the context of IAV-induced illness focused mostly on distinguishing IAV from non-IAV infections, symptomatic from asymptomatic IAV carriers, or low from high influenza vaccine responders (*14–20*). A notable exception was the recent description of blood transcriptomics data from a large cohort of subjects enrolled in the Mechanisms of Severe Acute Influenza Consortium (MOSAIC) study (*21*). In that report, severity of infection – measured in terms of need for mechanical ventilation – was associated with a weak transcriptional “viral response” signal and a strong transcriptional “bacterial response” (and activated-neutrophil) signal in blood in comparison with non-severe cases. However, these transcription patterns were also strongly associated with duration of illness and the authors emphasized the importance of timing in the interpretation of immune activity to IAV infection (*21*), which, for obvious reasons, can be hard to control for in natural human infection studies.

We aimed to develop a blood biomarker for early identification of individuals at risk of adverse disease outcome after influenza infection in a well-controlled mouse model of IAV infection, where a precise delineation of the evolution of the response associated with severity could be achieved. We utilized transcriptomic data from the lungs, the focal point of infection, and also data from blood post infection, to derive a transcriptional signature of lethality across various IAV and mouse strains. This early-response signature could distinguish mice at high risk of death with different influenza A virus strains. These results provide an impetus for seeking to translate this approach to human respiratory virus infection characterized by damaging inflammatory responses.

## Results

### An integrated tissue and blood multigene transcriptional signature of lethality

The first question was how to select a panel of candidate genes whose expression in blood could be used to predict lethality after IAV exposure. We focused on genes whose expression early after infection was associated with eventual lethal outcome, rather than with infection (*14, 16*). We reasoned that the focal point of infection, the lungs, where lethal processes unfold, would contain the relevant biological signal.

Several whole-genome transcriptomics datasets from mouse lung tissues after infection with IAV are publicly available, including from our own previous work (*13, 22, 23*). We utilized these resources (i.e., transcriptomic data from the mouse lungs at several time points after infection) to derive a gene signature of lethality from the lung across several IAV and mouse strains, followed by integration with blood data from influenza-infected mice and humans.

Briefly, to be considered for inclusion in the signature of lethality, on days 1 to 3 after infection genes had to be differentially expressed (DE), in comparison to PBS-treated animals, in the lungs of lethally infected mice but not DE in the lungs of non-lethally infected animals (see Fig. 1 and *Methods* for more details) (*13, 22, 23*). Furthermore, genes were included in the signature only if they were in at least one of the gene clusters associated with lethal, but not with non-lethal, IAV infections that we uncovered previously in the lungs (*13*).

**Figure 1:**
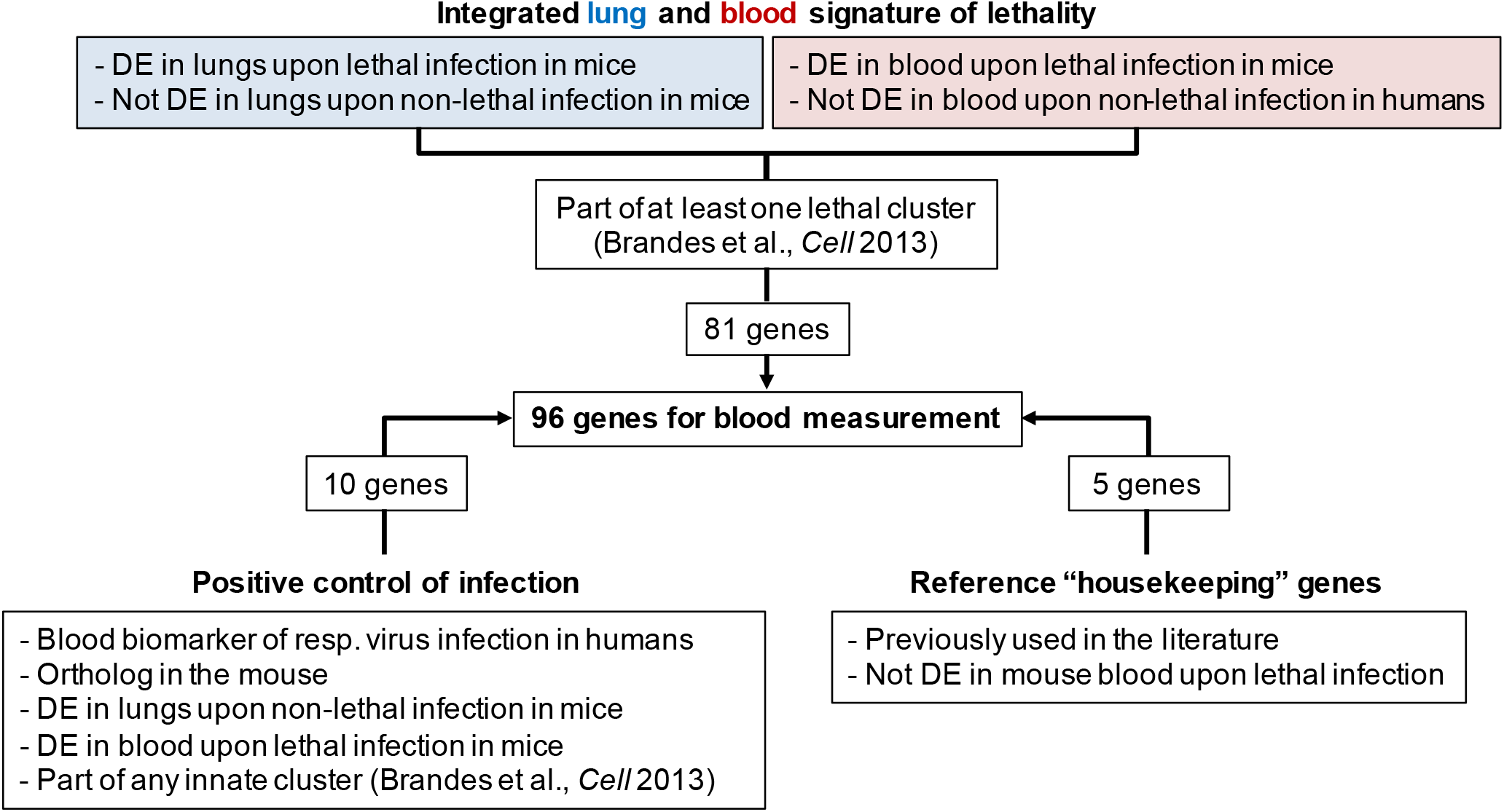
Criteria for selection of genes for the integrated tissue and blood signature of lethal influenza infection as well as positive control of infection and reference genes. DE: differentially expressed (Benjamini and Hochberg [BH] false discovery rate [FDR] = 0.1). The analysis included data from C57BL/6 and BALB/c mice infected with H1N1 PR8, H1N1 1918, H1N1 Tx91, and H5N1 VN1203, as well as from humans during the 2009 pandemic (*13, 22–24*).

The above procedure (the lung signature of lethal influenza infection) yielded 1,069 genes, which were enriched for several biological processes, including regulation of epithelial cell proliferation, cell adhesion, morphogenesis of an epithelium, regulation of vasculature development and regulation of immune system process (Benjamini and Hochberg [BH] false discovery rate [FDR] of Fisher's exact test of PANTHER overrepresentation: 2E-05, 3E-05, 1E-04, 1E-03 and 5E-02, respectively).

To refine this lung lethality signature with blood data, we only retained genes that were DE in the blood of mice two days after infection with a lethal dose of the highly pathogenic H1N1/PR8 IAV strain (Fig. S1) , and excluded genes that were DE in the blood of humans upon low pathogenicity infection (*24*). 81 genes passed these pre-selection filters. As a positive control for detection of IAV infection itself, independent of lethal outcome, 10 genes previously used as classifiers of respiratory virus infection in blood were selected (*14, 16*) as well as 5 reference “housekeeping” genes for normalization (*25*). Primers against 96 genes were designed for high throughput RT-qPCR (Table S1).

Since *(a)* our past work used non-lethal infection data to develop the specific set of lethality-associated gene clusters, *(b)* we excluded genes using human low pathogenicity data, and *(c)* we excluded genes DE in the lungs of non-lethally infected mice, the hope was that the candidate genes selected into the integrated tissue and blood signature would have better specificity for association with lethality.

### Training models for the classification of individual mice according to future outcome of disease

To determine whether the expression of these candidate genes in blood could be used as a starting point to train a machine learning model of lethality in mice after challenge with IAV, gene expression data were collected from RNA isolated from 50 μl of blood drawn from 34 animals four days after treatment with PBS or a range of lethal and sublethal influenza strain A/Puerto Rico/8/1934 H1N1 (PR8) doses (Fig. 2a-b). We used a statistical learning method known as elastic net and leave-one-out cross validation to assess whether predictive models could be built from our data (*26–28*).

**Figure 2:**
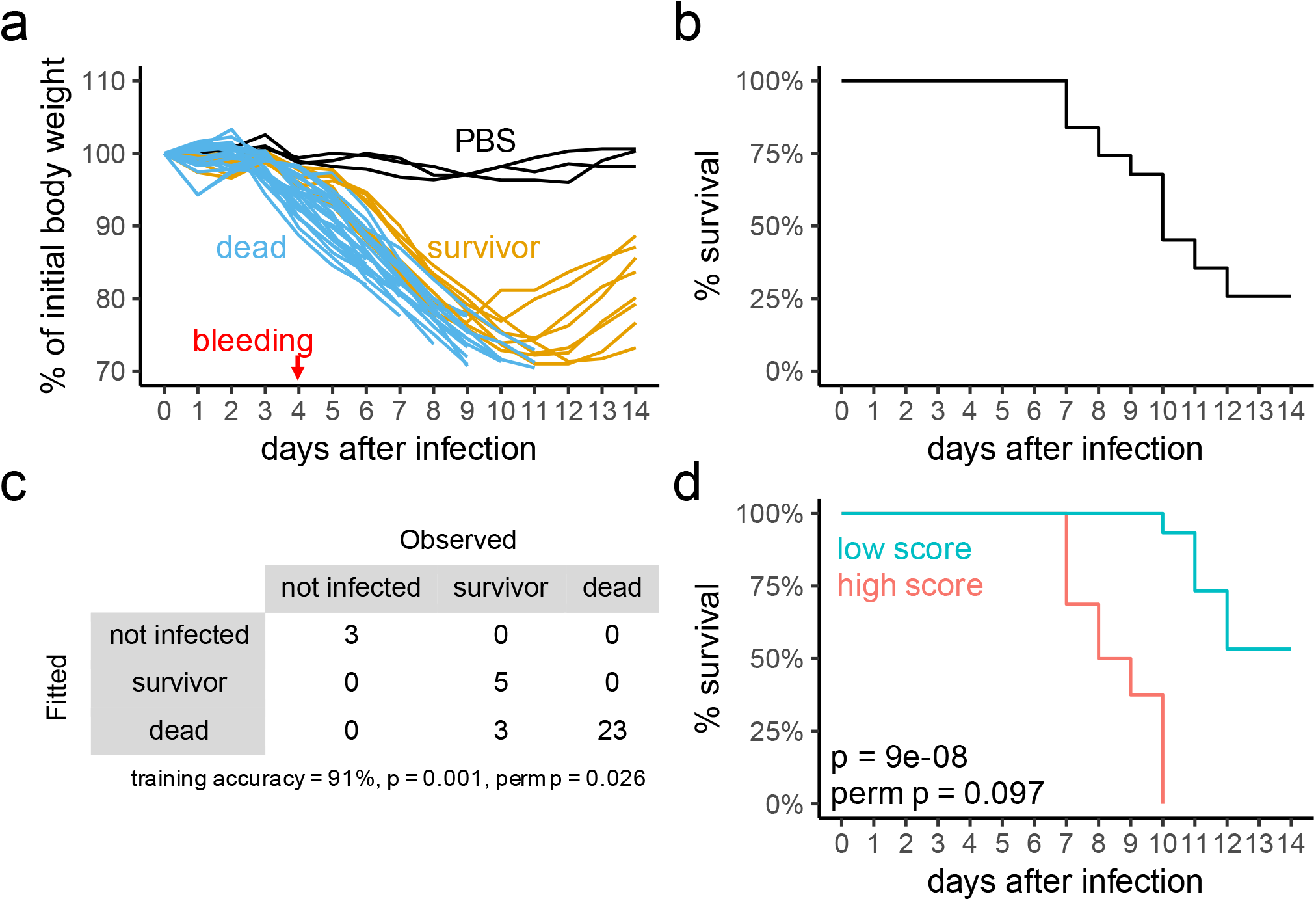
Training early multinomial classification and Cox survival models of lethal influenza A virus infection based on expression of candidate genes in whole peripheral blood. (a) Observed loss of body weight after treatment with PBS or infection with a range of PR8 doses. (b) Survival curve, excluding PBS-treated mice, after infection with range of PR8 doses. (c) Model fit after training for the multinomial model, showing confusion matrix for training error with observed and fitted events. P-value is from one-sided binomial test in comparison to “no information rate”. (d) Model fit after training for the Cox model, showing observed survival of mice with scores above (red) or below (blue) the median fitted score. P-value is from likelihood ratio test of Cox proportional hazards regression of survival on fitted relative risk, which did not violate proportional hazards assumption at alpha = 0.05 (cox.zph test). Perm p is the frequency of p-values in Monte Carlo permutations (perm) equal to or smaller than observed. N = 3 (PBS) and 31 (infected) in training cohort.

Briefly, survival (Fig. 2b) and gene expression data (Fig. S2) were fitted via elastic net-regularized multinomial logistic regression to generate a model to classify mice into three categories: 1) not infected, 2) surviving upon infection, or 3) dying upon infection. During training (i.e., model selection via cross-validation), the algorithm learned that 23 genes had expression values in blood that could be linearly combined to determine the probability of an individual animal being in one of the training categories (Fig. 2c and Fig. S3a). Some of the positive control genes were selected by the algorithm to help differentiate between infected and non-infected animals, and fewer to separate infected survivors from non-survivors (Fig. S3a), suggesting that the death-associated candidate genes were indeed enriched for detecting signals of lethality in blood.

### Training models to rank individual mice based on relative risk of early death

In the approach described above using logistic regression, the algorithm attempts to learn how the expression of each gene can be combined to discriminate mice in one category from another. In a hypothetical scenario of resource prioritization, however, one might also be interested in estimating the time to a relevant clinical event – death, in this case. This can be achieved with Cox proportional hazards regression, where time to event is taken into consideration and the relative risk of death for each subject can be derived.

Hence, to examine whether the expression of lethal candidate genes in blood would distinguish mice at high risk of early death after challenge, we combined survival and gene expression data from the 81 lethal candidate genes in Cox regression regularized via the elastic net. During training (i.e., model selection via cross-validation), the algorithm learned that 11 genes had expression values in blood that could be linearly combined to generate a scoring system of infected animals as a function of the day of death (Fig. 2d and Fig. S3b). Training with positive control of infection genes failed to yield predictive models (leave one out cross-validation p = 0.15, vs. p = 0.009 when lethal candidates were used in leave one out cross-validation [p values from likelihood ratio test of Cox proportional hazards regression of survival on fitted-relative risk]), underlining the ability of the affected tissue signature to capture information associated with lethality.

In both the multinomial and the lethal Cox models, high levels of expression of genes associated with monocytes and neutrophils, together with low levels of transcripts associated with lymphocytes, indicated high risk of death (Fig. S3), consistent with previously described analyses of the immune response to IAV infection (*10, 12, 13*) and also recent data from COVID-19 patients (*29*). *Art4*, the gene most positively associated with lethality in both models, is highly expressed in hematopoietic stem cells and immature lymphocytes according to the Immgen database (*30*), indicating a potential association of lethality with dysregulated hematopoiesis and release of immature cells into circulation.

### Prediction of lethality with independent cohorts of mice

An important aspect of machine learning-derived models is whether their performance is generalizable, which means whether they perform well on unseen test data that has not been used in training. Here, the models were tested for their ability to predict lethality based on gene expression data from independent cohorts of mice that were not available for training. Performance was tested in three different ways: 1) on mice infected with high doses of either a low or a high pathogenicity influenza strain (i.e., non-lethal high dose strain A/Texas/36/91 H1N1 (Tx91) vs. lethal high dose PR8); 2) by training our models on this second dataset (low vs. high pathogenicity data) while testing on our first dataset described above from infection with different doses of PR8; and 3) by testing on the scenario of infection at one lethal dose 50 (LD_50_), where mice are challenged with virtually the same viral dose but only half of them survives the infection.

In the first round of validation, an independent cohort of 14 mice bled on day two after treatment with PBS or with a high dose of either Tx91 or PR8 provided the data (Fig. 3a-b). Although the test set was from an earlier time point of infection and included one virus strain and one dose not used in training, both the multinomial and the lethal Cox models showed good predictive accuracy (Fig. 3c-d). In the second round of validation, after reversing the roles of each dataset (i.e., training on the set with two different IAV strains on day two of infection and testing on the set with five different doses of PR8 on day four of infection), the multinomial model did not perform as well as the lethal Cox model, which showed good accuracy for predicting outcome (Fig. S4a-b).

**Figure 3:**
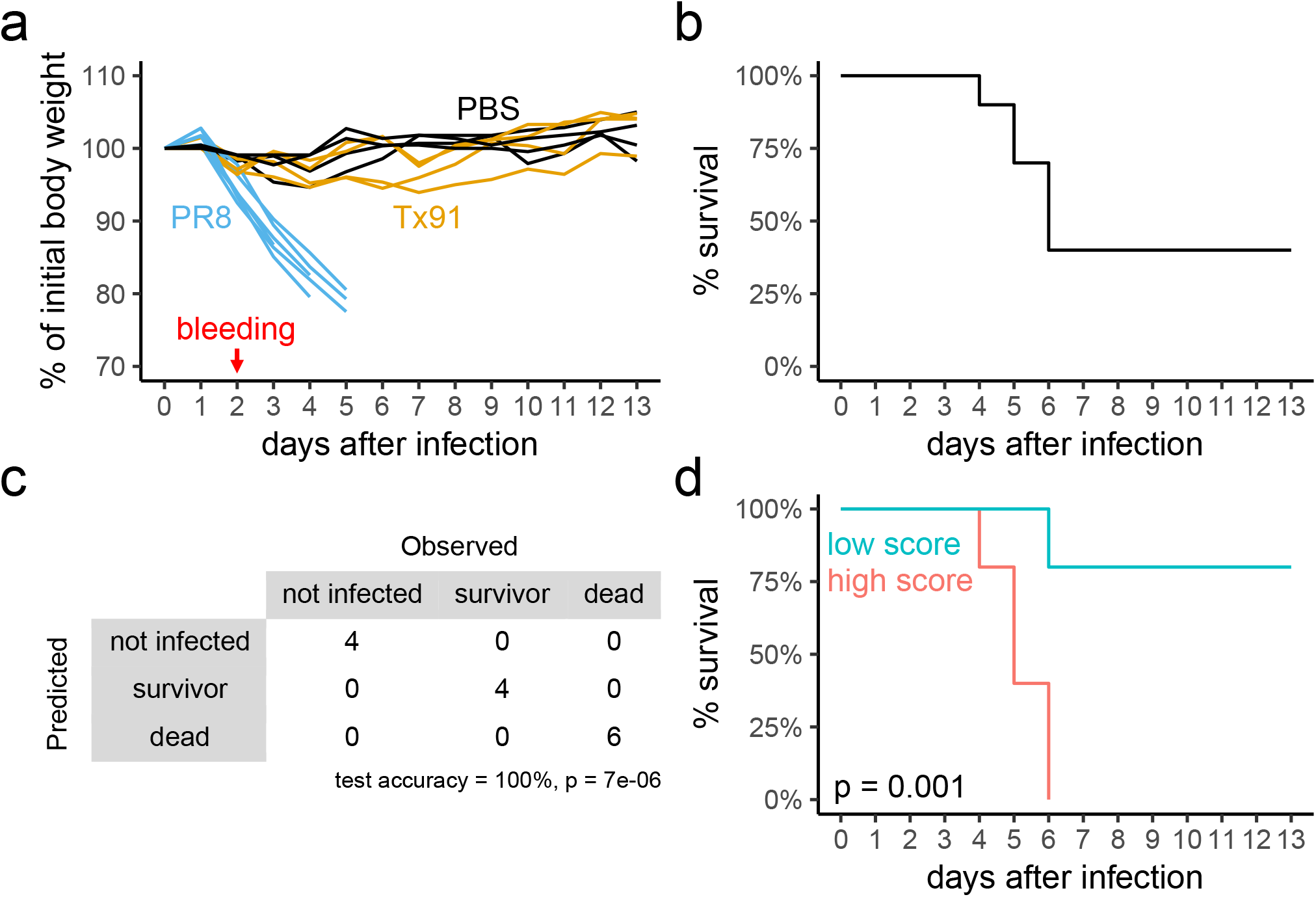
Testing early models of lethal influenza A virus infection on independent cohorts of mice. (a) Observed loss of body weight after treatment with PBS or infection with high doses of Tx91 or PR8. (b) Survival curve, excluding PBS-treated mice, after infection with high doses of Tx91 or PR8. N = 4 (PBS), 4 (Tx91) and 6 (lethal PR8). (c) Model performance on this independent cohort of mice for the multinomial model, showing confusion matrix for prediction error with observed and predicted events. P-values are from one-sided a binomial test in comparison to “no information rate”. (d) Model performance on this independent cohort of mice for the Cox model, showing observed survival of mice with scores above (red) or below (blue) the median predicted score. P-values are from likelihood ratio test of Cox proportional hazards regression of survival on predicted relative risk, which did not violate proportional hazards assumption at alpha = 0.05 (cox.zph test).

Considering only the lethal Cox model, for our third round of validation we turned our attention to the challenging scenario of predicting lethality among genetically identical, sex and age matched, cohoused mice given the same infectious bolus at an LD_50_ of PR8 (N = 24 mice). In this setting, typically about half of the mice succumb while the other half survives viral exposure. Importantly, it is not known mechanistically what drives this outcome dichotomy, since measuring putative candidate factors such as lung viral titer and immune infiltration requires sacrificing the mice and, thus, precludes assessment of the actual outcome of disease. To us, the most obvious candidate mechanism was that differences in initial infectious bolus effectively received by each animal during the infection procedure would determine the dichotomy. In that case, our lethal model should be able to predict the outcome of infection very early on, and thus help shed light on the biological mechanisms of life or death under these particular conditions.

### Lack of signs of infection early on at 1 LD_50_ PR8

Incubation period is the time that transpires from the moment of exposure to a pathogenic agent to when the host starts showing signs of infection. During this period, a virus needs to arrive at accessible tissues and bind to receptors to enter and replicate in susceptible cells. From infected cells, new infectious viral particles are released, and the cycle starts again, with viral titers temporarily following an exponential growth curve at least while virus replication is unperturbed by the host immune system. Naturally, the time to trigger effective host immunity is influenced by viral virulence, replication capacity and initial infectious dose.

In our hands, no systemic signs of infection could be detected in blood using positive control of infection genes two days after challenge with 1 LD_50_ PR8 (Fig. S5a-c). In contrast, mice on day two of infection in our previous experiment – i.e., challenged with high dose low pathogenicity Tx91 or high dose PR8 – showed changes in gene expression in several infection associated positive control genes (Fig. S5c). Considering that only a very small number of PR8 virions is required to induce pathogenic lung disease, these results indicate that on day two of 1 LD_50_ PR8 infection the virus had not yet reached high enough levels in the respiratory tree for the early anti-viral response to become detectable in blood, and, thus, our model was not able to detect any signs of lethality in the blood of mice on day two post 1 LD_50_ PR8 infection.

### Correct prediction of risk of death at the group level but lack of resolution for within group inter-individual distinction on day 4 after 1 LD_50_ PR8 infection

By day four, however, DE of all 9 positive control of infection genes could be detected in the blood of animals infected with 1 LD_50_ PR8 (Fig. S5c), likely reflecting the spread of PR8 in the lungs and evolving host immunity. In addition, on day four, our lethal model correctly placed the animals at an intermediate level of risk of death – higher than mice infected with low pathogenicity Tx91, where no mice were at any risk of death, and lower than high dose lethal PR8, where every animal would eventually succumb (Fig. S5d). However, on day four, our model was unable to predict accurately which of the individual mice within the 1 LD_50_ group would survive and which would die (Fig. S5e). Retraining our model on 1 LD_50_ blood gene expression data also failed, as did predictions based on loss of body weight up to day four, suggesting that the fate of these mice might be indistinguishable at this early point after infection.

### Specific time window for within group, inter-individual prediction of lethality upon 1 LD_50_ PR8 infection

In an attempt to further characterize the infection dynamics, we devised a longitudinal experiment in which ethically small samples of blood (~10 μl) were taken daily from the same animals in a different cohort of 16 mice during days four to eight of infection with 1 LD_50_ PR8 (Fig. 4a), followed by RNA isolation, high-throughput RT-qPCR, and testing based on the existing lethality model (i.e., without retraining). The model had statistically significant power to distinguish within group, inter-individual differences in relative risk of death after 1 LD_50_ challenge on days five and six, but not on days four, seven or eight (Fig. 4b). These results suggest a specific time window within which processes associated with LD_50_ lethality is reflected by our signature in blood. While the lack of accuracy later in the course of infection (i.e., days seven and eight) was likely due to the fact that the model was trained for early detection of lethality, before adaptive immune cells fully developed and reached the circulation, these data suggest that PAMPs and DAMPs eventually reach different levels in the lungs of different mice, impacting their blood cell composition and lethal score, which can be used for prediction of lethality of individual animals even in the challenging scenario of experimental infection with an 1 LD_50_ IAV dose.

**Figure 4:**
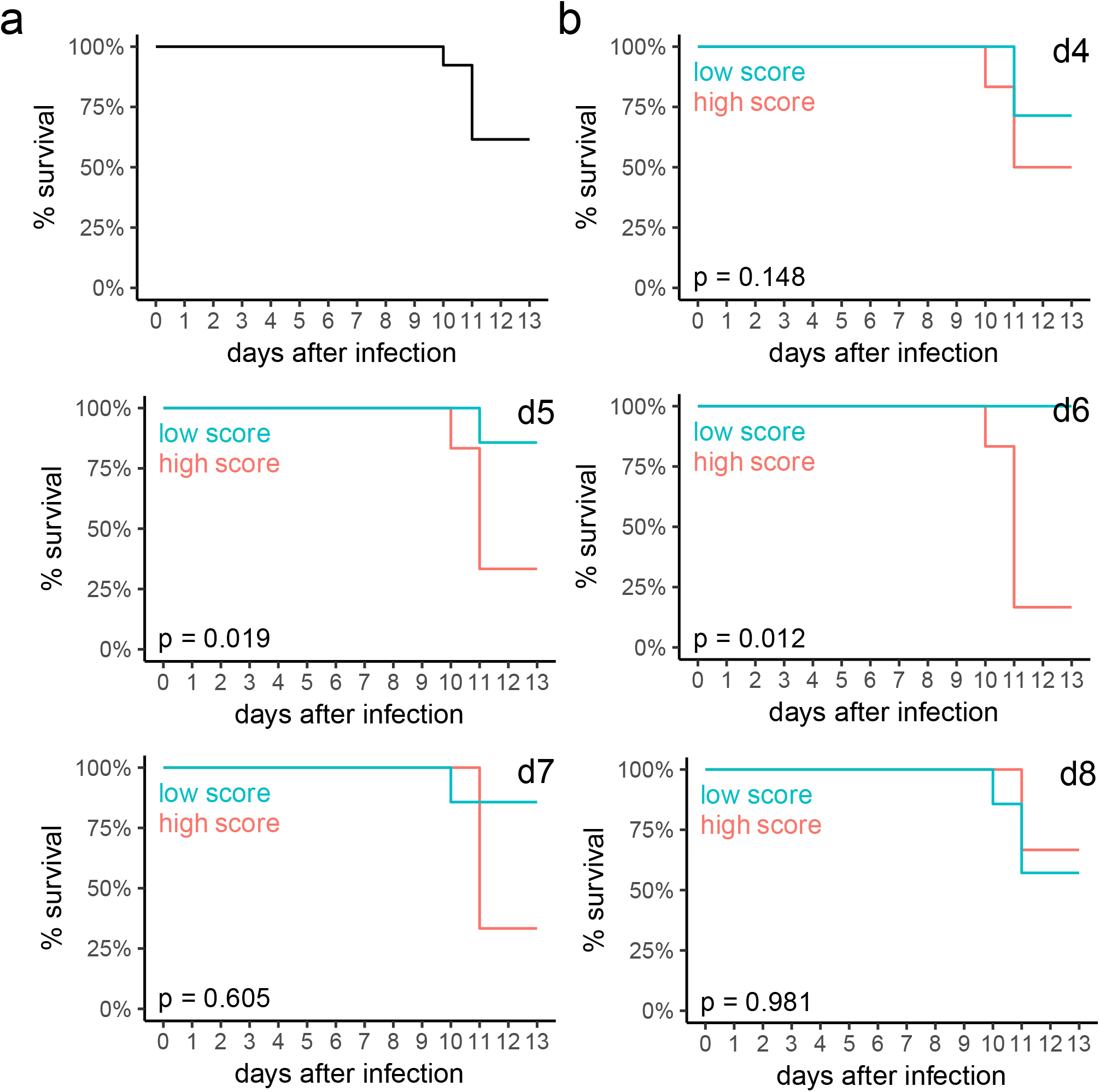
Prediction of outcome of disease in lethal influenza A virus infection on different days after challenge with 1 LD_50_ PR8. (a and b) Similar to Fig. 3d, but the results of predicting survival on different days of infection with 1 LD_50_ PR8 are shown. P-values are from likelihood ratio test of Cox proportional hazards regression of survival on predicted relative risk, which did not violate proportional hazards assumption at alpha = 0.05 (cox.zph test). P-values were adjusted using BH. N = 16.

## Discussion

Prediction tools for infectious disease outcomes would enable evidence-based treatment decisions and resource allocation, in particular during pandemics. We show that a transcriptional signature predictive of lethality can be developed via machine learning in a mouse model of influenza infection early after exposure, from as little as 10 μl of blood. This was achieved by integrating tissue and blood signatures of lethal influenza infection. The model also revealed a specific time window for prediction of lethality in the challenging scenario of 1 LD_50_ PR8 infection.

Earlier in the course of infection, our model tended to place mice infected with 1 LD_50_ PR8 at an intermediate risk level in comparison to mice infected with high dose PR8 or Tx91, suggesting that our model can delineate effects associated with infection severity. In the scenario of LD_50_ PR8 infection of inbred and cohoused littermates with virtually the same infectious bolus, our model revealed a later split (~days 5 and 6) between animals who would eventually die or survive.

These intriguing results raise the possibility that differences in initial infectious bolus were not the primary determinants of life or death of mice infected with 1 LD_50_. Rather, these mice seem to behave like one group undergoing similar responses until they reach the boundaries of a tipping point, with survival or death being the result of biological variation around that tipping point (schematic model in Fig. S5f). The time when these differences were observed (i.e., days 5 and 6) overlaps with the early effector phase of adaptive immunity.

Since adaptive immunity, in particular cytotoxic T cells, plays an important role in reducing viral load and, hence, in stopping the feedforward stimulation of the damaging innate response (*13, 31–43*), the timing when predictions can be made suggests that early inter-individual differences in the adaptive response may make a critical contribution to differences in the outcome of disease. Although the average T cell response to infection is remarkably efficient and constant (*44–48*), formation of the naïve repertoire of antigen-specific T cells is a semi-stochastic process that varies in each host, resulting in a range of precursor frequencies (*49–51*) and, presumably, leading to a spread among the mice in the kinetics of reaching an adequate effector T cell number for effective viral clearance.

Therefore, the timing of the split between survivors and non-survivors in the 1 LD_50_ scenario potentially reflects differential interference with virus production during the early phase of the effector T cell response, which may be the determinant of life or death at the tipping point of infection. However, a cause/effect relationship is difficult to test directly due to the lack of tools to accurately examine, for example, virus titer, lung infiltration by antigen-specific cells, or the size of their precursor population in live mice, because these would require terminal experiments that would preclude determination of which animals would eventually die several days later due to the actual infection. Nonetheless, preliminary experiments adding gene probes to our panel that report activated CD8+ T cell abundance in the blood samples suggest that there is a relationship between the strength of these lymphocyte-related signals in day 5 or 6 samples and survival after infection. Further tests along these lines are planned to refine the signature and relate these blood findings in a large cohort of infected animals to effector function and viral abundance in the lungs as directly assessed by multiplex tissue imaging.

Our experiments have several limitations, such as the lack of validation of the models on human IAV infection data. Unfortunately, to our knowledge the only report to contain both whole blood whole-genome transcriptomics and disease severity data that includes severe human IAV infection, contained data from only three severe cases beginning five days after onset of symptoms or earlier (*21*). Beside the fact that such low number of early cases preclude attempts to validate our models, one also needs to take into consideration the incubation time – the time from exposure until development of symptoms – when considering the temporal evolution of infection.

The models presented here were developed for prediction very early after exposure, and, while animal models of infection can be used to clearly delineate the evolution of the immune response along its temporal component, it remains to be determined which time window post exposure in mice most closely translates to time after onset of symptoms in human infection. Experimental IAV challenge in humans with relatively mild strains indicates a peak of symptoms (mostly runny nose and sore throat but no need for mechanical ventilation) around day 3-4 after exposure (*52*), but it is not clear how long it takes, on average, for a symptomatic carrier to seek medical treatment, in particular in case of severe disease with highly virulent and/or highly pathogenic strains. Such questions would need to be addressed to enhance preparedness for the next pandemic and before predictive models developed in animal experiments could be realistically deployed to human populations, including with respect to the ongoing SARS-CoV2 pandemic.

Although mice and humans have substantial differences, including different distributions of influenza viral receptors in the respiratory tree, which is crucial to create an inter-species barrier for transmission, the immunobiology of highly pathogenic influenza infection, once transmission has been established, is quite similar across the several species that have been studied with regard to strong innate immune activation (*7, 53, 54*). Given the similarities between this animal model of lethal viral infection and the course of human infections with certain respiratory virus strains (*53–59*), our findings provide impetus in the development of prognostic tools for managing patients with severe viral respiratory infection as well as clues that may guide development of interventions and timing for their administration.

## Acknowledgments

New expression profiling by high throughput sequencing data reported here are available on GEO with accession number <u>GSE124404</u>. We would like to thank Emily Condiff, Dr. Antonio P. Baptista, Dr. Stefan Uderhardt and Dr. Sonja M. Best for critically reviewing the manuscript as well as members of the Laboratory of Immune Systems Biology at NIAID for fruitful discussions, in particular the members of the Lymphocyte Biology Section. This study used the Office of Cyber Infrastructure and Computational Biology (OCICB) High Performance Computing (HPC) cluster at the National Institute of Allergy and Infectious Diseases (NIAID), Bethesda, MD. This work was supported by the Intramural Research Program of NIAID, NIH. Luis M. Franco is supported by the Intramural Research Programs of NIAID and NIAMS, NIH.

## Supplemental Figures

**Figure S1:**
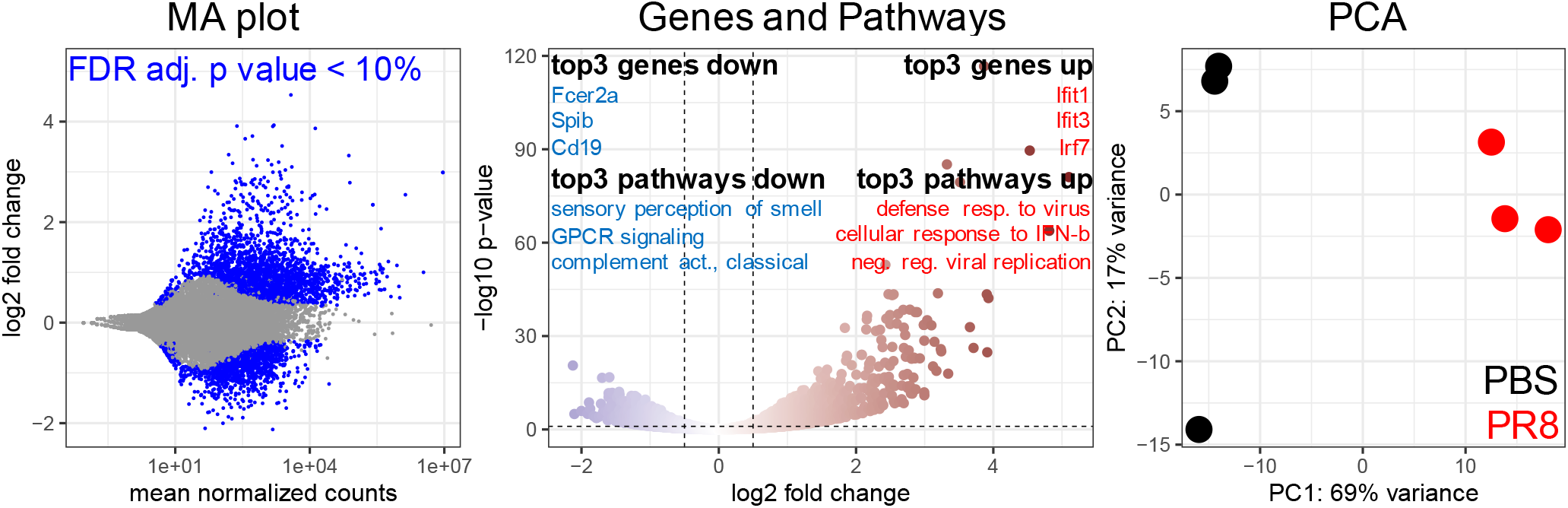
Whole genome transcriptomics results from the blood of mice treated with PBS or infected with high dose PR8. DESeq2 analysis of blood RNA-seq data comparing infected and non-infected mice where genes were considered DE between groups at BH-FDR < 0.1. Left hand panel: MA plot with genes colored according to DESeq2 results (DE genes: blue, non-DE gene: gray). Middle panel: Volcano plot with genes colored according to log2 fold change, as well as results from PANTHER overrepresentation test using DE genes (the top 3 most significantly up and down regulated genes and pathways in the blood of infected animals are highlighted). Horizontal dashed line: BH-FDR = 0.1. Vertical dashed line: log2 fold change = −0.5 and 0.5. PANTHER: BH-FDR was smaller than 0.05 for all top 3 up and down regulated pathways in Fisher’s exact test. Right hand panel: PCA results with samples colored according to treatment (PBS: black, PR8: red). N = 3 (PBS) and 3 (lethal PR8).

**Figure S2:**
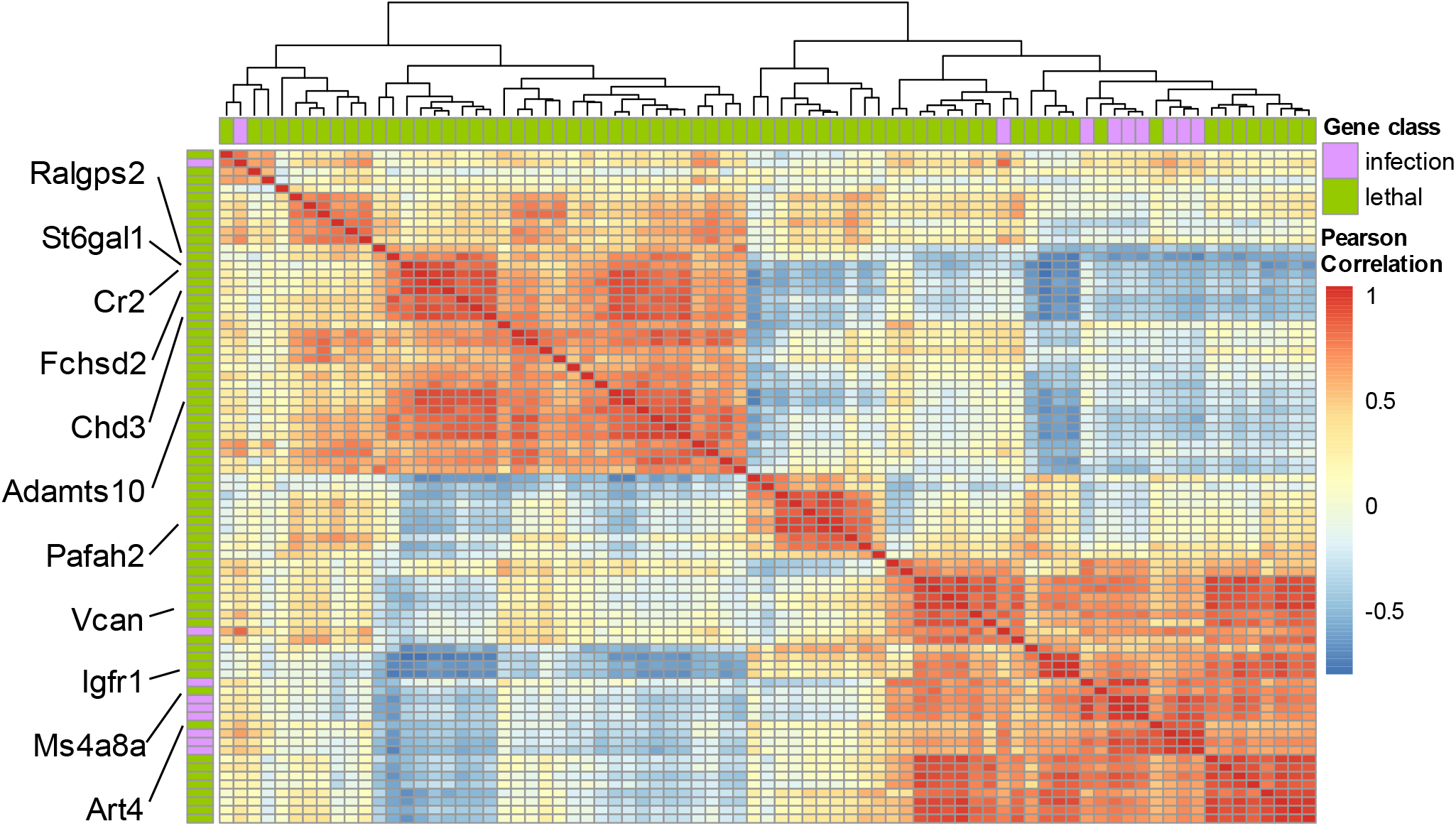
Expression of target candidate lethal genes and positive control of infection genes in blood of IAV infected mice. Pearson correlation heatmap of blood gene expression data on day four after infection with a range of PR8 doses, including positive control of infection (magenta) and lethal candidate (green) genes. 11 genes that are part of the lethal Cox model are highlighted (Fig. S3b).

**Figure S3:**
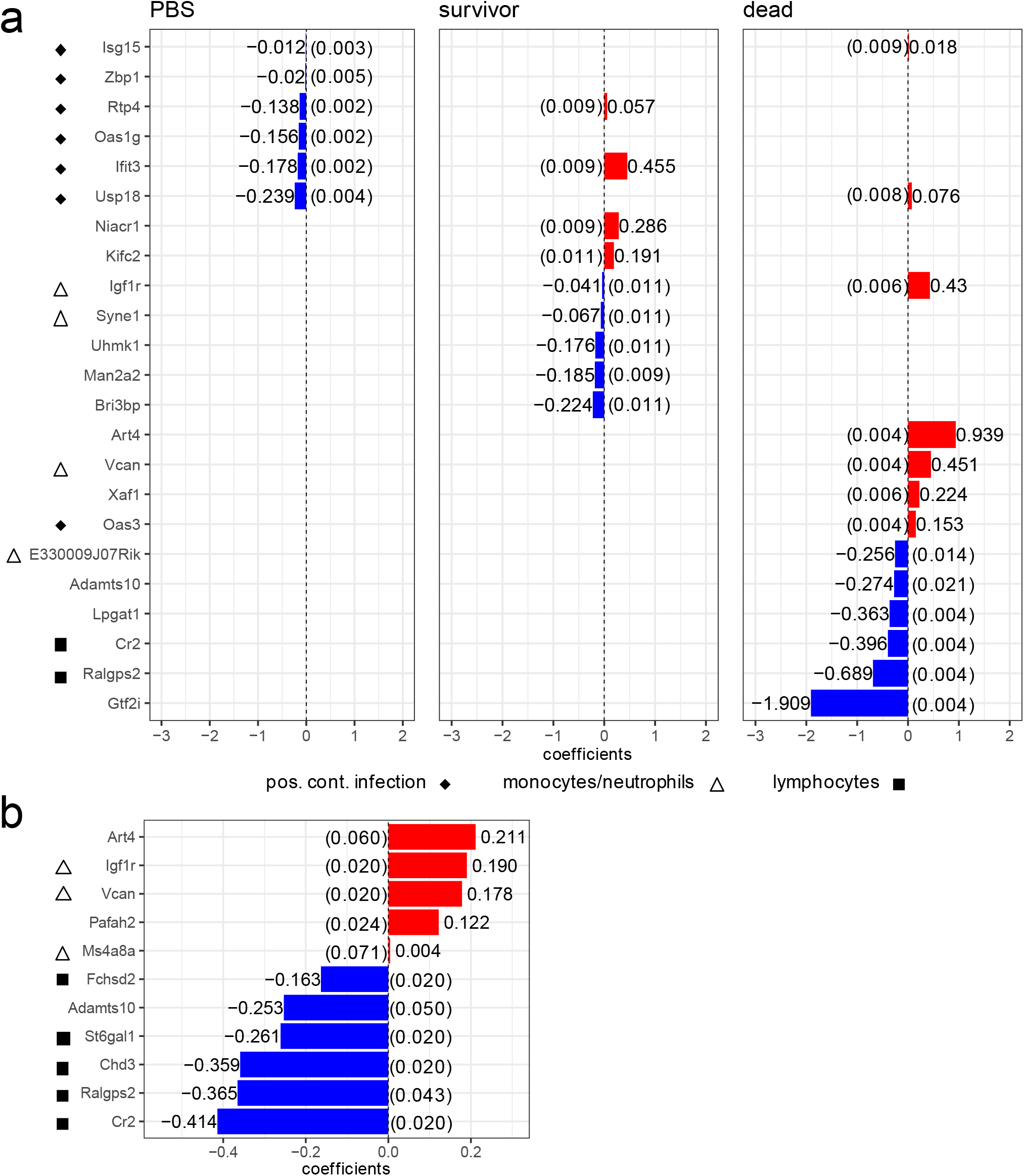
Multinomial model and Cox model coefficients. (a) Inverted coefficients of classification model of lethal influenza (coefficients were inverted for visualization, since lower deltaCt values in RT-qPCR mean higher gene expression levels). Each box represents a category (not infected mice, survivor upon infection, dead upon infection) and bars show the inverted coefficient given to each gene to calculate the probability of a mouse belonging to a respective category. (b) Inverted coefficients of Cox model of lethal influenza (as above, inverted for visualization). Symbols to the left of gene names indicate whether genes were among positive control of infection genes, or, if not, which cell types had highest expression levels of the given gene according to the ImmGen database. (a and b) Numbers in parentheses are empirical p-values from Monte Carlo permutations (frequency of coefficients in MC permutations that were equal to or larger in magnitude than observed [one-sided to ensure agreement in direction of association]) after BH-adjustment.

**Figure S4:**
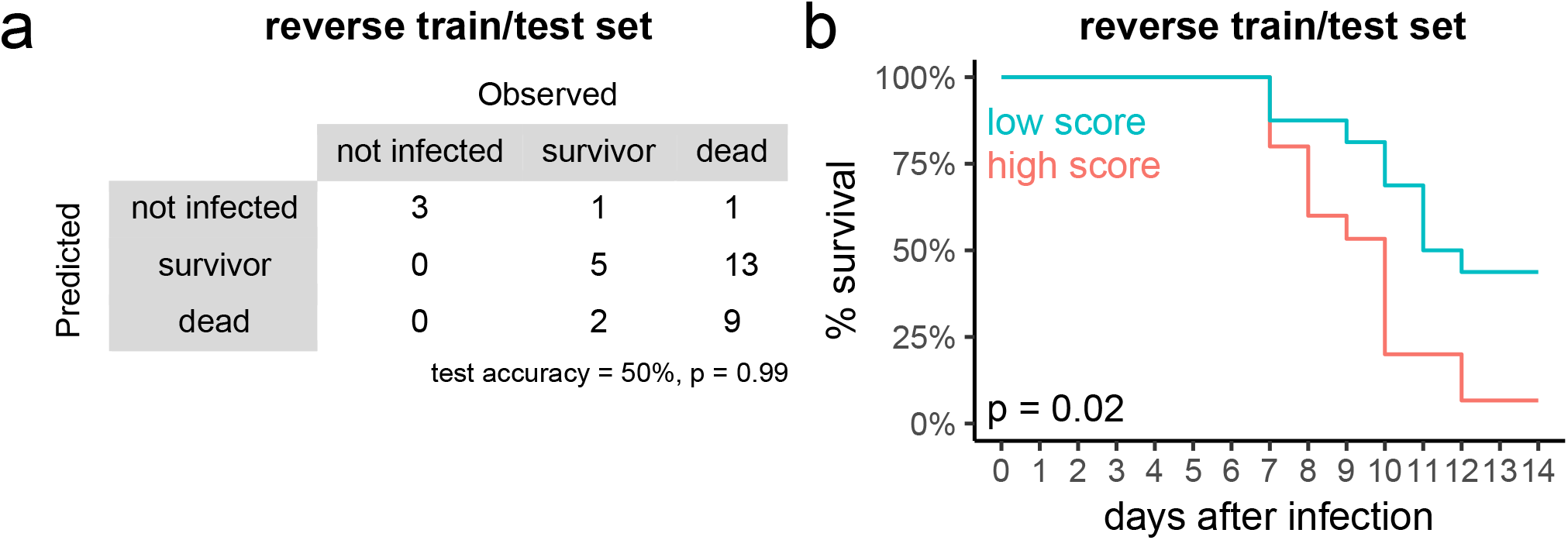
Validation with reversed training and test sets. Model performance when reversing the roles of the datasets (i.e., training on the high dose Tx91/PR8 cohort and testing on the PR8 dose range cohort), showing either confusion matrix for prediction error with observed and predicted events (a) or observed survival of mice with scores above (red) or below (blue) the median predicted score (b). In (a), p-values are from one-sided a binomial test in comparison to “no information rate”. In (b), p-values are from likelihood ratio test of Cox proportional hazards regression of survival on predicted relative risk, which did not violate proportional hazards assumption at alpha = 0.05 (cox.zph test).

**Figure S5:**
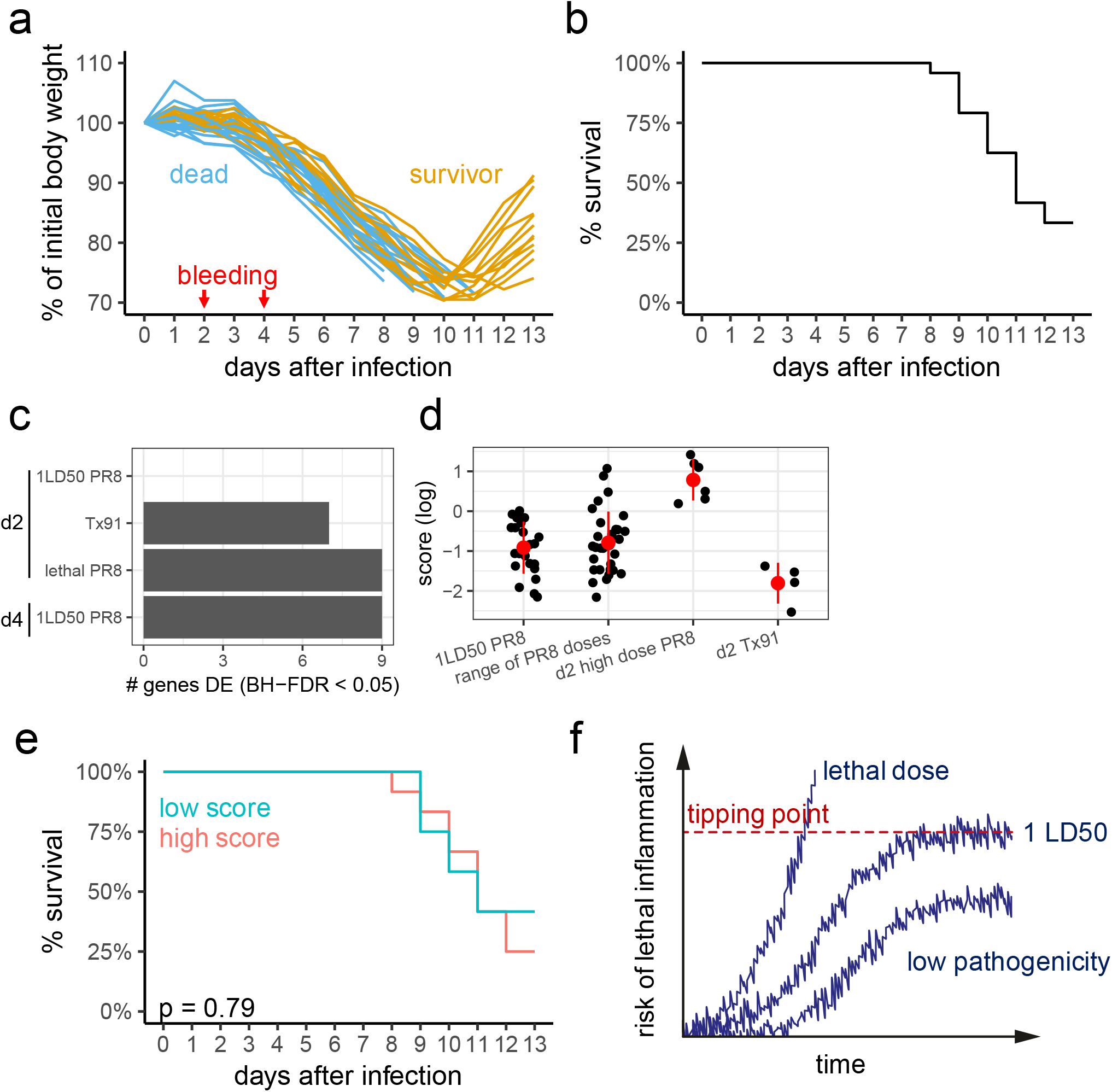
Test error on data from the blood of mice early after infection with 1 LD_50_ PR8. (a/b/e) Similar to Fig. 3a/b/d, but the results of testing the model on data from 1 LD_50_ infection experiments are shown. In (e), p-value is from likelihood ratio test of Cox proportional hazards regression of survival on predicted relative risk (did not violate proportional hazards assumption at alpha = 0.05 [cox.zph test]). (c) Number of positive control of infection genes DE in the blood of mice on days two and four after infection with high dose PR8, Tx91 or 1 LD_50_ PR8. Expression levels of positive control of infection genes were compared with PBS-treated mice (BH-FDR < 0.05 [Mann-Whitney test] for a gene to be considered DE). (d) Predicted relative risk (log) for each mouse is shown for comparison, with the red dot and red line representing mean and standard deviation in each group. (f) Schematic model of risk of lethal inflammation rising with time early after infection with different doses of and strains IAV. N = 31 in a/b (all infected mice), and 24 from c to e (mice with blood RNA samples that passed QC).

## Methods

### Data visualization and analysis

Data visualization and analysis was performed in R 4.0.2 and RStudio (*60*), using packages downloaded from CRAN and Bioconductor 3.11 (*61*). Data were handled, plotted and visualized with the foreach (*62*), dplyr (*63*), ggfortify (*64*), ggplot2 (*65*) and gridExtra (*66*) packages.

### Mouse model of infection

Male C57BL/6J-CD45a(Ly5a) mice were obtained from Taconic Laboratories (line 8478). Animals aged 8 to 12 weeks were used. Mice were maintained in specific-pathogen-free conditions at an Association for Assessment and Accreditation of Laboratory Animal Care-accredited animal facility at NIAID and were used under study protocols (LSB-1E, LSB-4E, LISB-1E and LISB-4E) approved by the NIAID Animal Care and Use Committee (National Institutes of Health). Influenza H1N1 A/PR/8/34 and A/Tx/91 propagation and infection procedures were described before (*13, 67*). PR8 doses used: 0.5 LD_50_, 1 LD_50_, 2 LD_50_, 4 LD_50_ and 8 LD_50_ (model training experiment) and 100 LD_50_ (RNA-seq and model validation experiments). Tx91 dose used: 1 million PFU (model validation experiment). Mice were excluded if they sneezed during the infection procedure and failed to lose body weight in a similar manner as other mice in the same treatment groups.

### Global gene expression analysis

For RNA-seq, blood was collected postmortem via heart puncture on day two after infection (n = 3 mice/treatment) and kept at −20 °C in RNAlater (Thermo Fisher Scientific). RNA was extracted using the Mouse RiboPure Blood RNA Isolation Kit and globin RNA depleted using the GLOBINclear Kit, mouse/rat (Thermo Fisher Scientific). Quality and amount of RNA were determined using the Qubit RNA BR Assay Kit with Qubit 2.0 (Thermo Fisher Scientific) and Agilent RNA 6000 Nano Kit on the Agilent 2100 Bioanalyzer. Only samples with RIN score above 7 were further analyzed. Libraries were prepared using NEBNext Ultra Directional RNA Library Prep Kit for Illumina, NEBNext^®^ Multiplex Oligos for Illumina^®^ (Index Primers Set 1 and 2) and NEBNext Poly(A) mRNA Magnetic Isolation Module (New England Biolabs). Library quantification was performed by real-time PCR using the KAPA Library Quantification Kit - Illumina Platforms - Complete kit (Universal) and KAPA Library Amplification Kit - Real-time PCR library amplification, with fluorescent standards (KAPA Biosystems). Libraries were sequenced twice on the Illumina Nextseq platform (Illumina) using V1 reagents to a total read depth of 60 million paired-end, 75 base-pair reads. Reads were mapped to the *M. musculus* UCSC mm9 genome assembly using Bowtie2 (*68*). Counting of reads mapping to each gene was performed in R using *featureCounts* from the R package Subread. The count matrix was processed with DESeq2 (*69*). Genes were considered differentially expressed in infected mice in comparison to PBS-treated animals at BH-FDR of 0.1. PANTHER overrepresentation test (*70*) was performed online (http://www.geneontology.org; released 20181010) for GO biological processes in *Mus musculus* using Fisher’s exact test and BH adjustment.

### Integrated tissue and blood signature of lethal influenza infection

A list of candidate genes for the affected lung tissue signature of lethal influenza infection, as well as respiratory virus infection positive controls and reference genes was created by checking for overlap between the gene lists that were either downloaded from the original publications or created using the GEO data with the GEO2R tool (BH-FDR = 0.1; see text and Fig. 1 for details) (*13, 22–24*). When Entrez gene IDs were not available, the Mouse Genome Informatics database (Jackson) was used to find current and old gene symbols as well as synonyms. A final list with selected genes and primers is shown in Table S1.

### Targeted gene expression analysis

For model training and validation experiments, around 50 μl of blood were collected from the submandibular vein on indicated time points and kept at −70 °C in TRIzol LS (Thermo Fisher Scientific). Groups of mice (3-4 mice/dose) were treated in two independent infection experiments for training and other independent experiments for validation, totaling 6-8 mice per dose. In addition, the first batch of mice infected with 1 LD_50_ PR8 had 34 animals from two independent experiments. For the 1 LD_50_ longitudinal experiment, 20 mice were infected and 10 μl of blood were collected from the tail vein on indicated time points and kept at −70 °C in TRIzol LS. RNA was extracted using the Direct-zol RNA Microprep Kit (Zymo Research). Quality and amount of RNA were determined using the Qubit RNA BR Assay Kit with Qubit 2.0 (Thermo Fisher Scientific) and Agilent RNA 6000 Nano Kit on the Agilent 2100 Bioanalyzer. Only samples with RIN score above 7 were further analyzed. Primer design and gene expression analysis were performed with the 96.96 IFC using Delta Gene Assays software (D3 Assay Design) and protocols for pre-amplified samples for use with Biomark HD, following the manufacturer’s instructions (Fluidigm) as described previously (*71*). To generate a standard curve for assessment of primer performance, samples from infected and non-infected animals were pooled and applied in a standard curve in duplicates (two no-sample controls were also included). Initial quality control was performed on the Biomark HD using Real-Time PCR Analysis v4.1.3 (Fluidigm), with automatic threshold generation. Genes with primers unable to generate high quality results across the range of dilutions of the standard curve (R^2^ > 0.85 in Ct value vs. dilution plots) or generating more than one peak in the melting curve were excluded from further analysis.

### Machine learning: training

Data was imported to R and normalized with HTqPCR (*72*). Normalization was based on the standard delta Ct method (subtraction of the mean of the reference “housekeeping” genes from all other values). In the machine learning algorithm, training was performed with glmnet (*26–28*) using leave-one-out cross-validation to tune the regularization parameter lambda (alpha = 0.5, family = “multinomial” for classification model [PBS vs. survivor vs. dead], family = “cox” for scoring model). Importantly, training of the scoring model was performed excluding PBS-treated mice and positive control genes. On the other hand, training of the classification model did include PBS-treated animals and positive control genes. perm p (empirical p value from Monte Carlo permutations) of the training error (estimating how often a result at least as good as the observed training error is expected by chance in a random, permuted training set) and perm p of the feature coefficients (estimating how often a coefficient at least as large in magnitude and in the same direction [positive vs. negative] as the observed coefficient is expected by chance in a random, permuted training set) were estimated using 1000 Monte Carlo permutations (*73*) either by shuffling the rows of the survival matrix, where time and status are the only two columns (Cox models), or by shuffling the elements of the vector of classes (multinomial regression).

### Machine learning: testing (cross-validation and validation on independent data)

The test error of the scoring model was estimated both using leave-one-out cross-validation within the training cohort (also known as nested cross-validation, in which the leave-one-out procedure is used *(a)* in a sub-cohort of the training samples to tune the hyperparameter lambda, followed by predicting on the one sample not included in that sub-cohort, and *(b)* repeated for every sample in the training cohort) and on an independent cohort of mice. A Cox proportional hazards regression model was fitted to the predicted score (i.e., the relative risk derived using the predict function of the glmnet package [type = response]) of each mouse using the survival package and checked that they did not violate the proportional hazards assumption at alpha = 0.05 using the cox.zph function also in the survival package (*74*). Importantly, the low number of mice per category precluded the use of cross-validation to estimate the test error of the classification model, which was done only using the independent cohort of mice.

## Supplemental Table

**Table S1:**
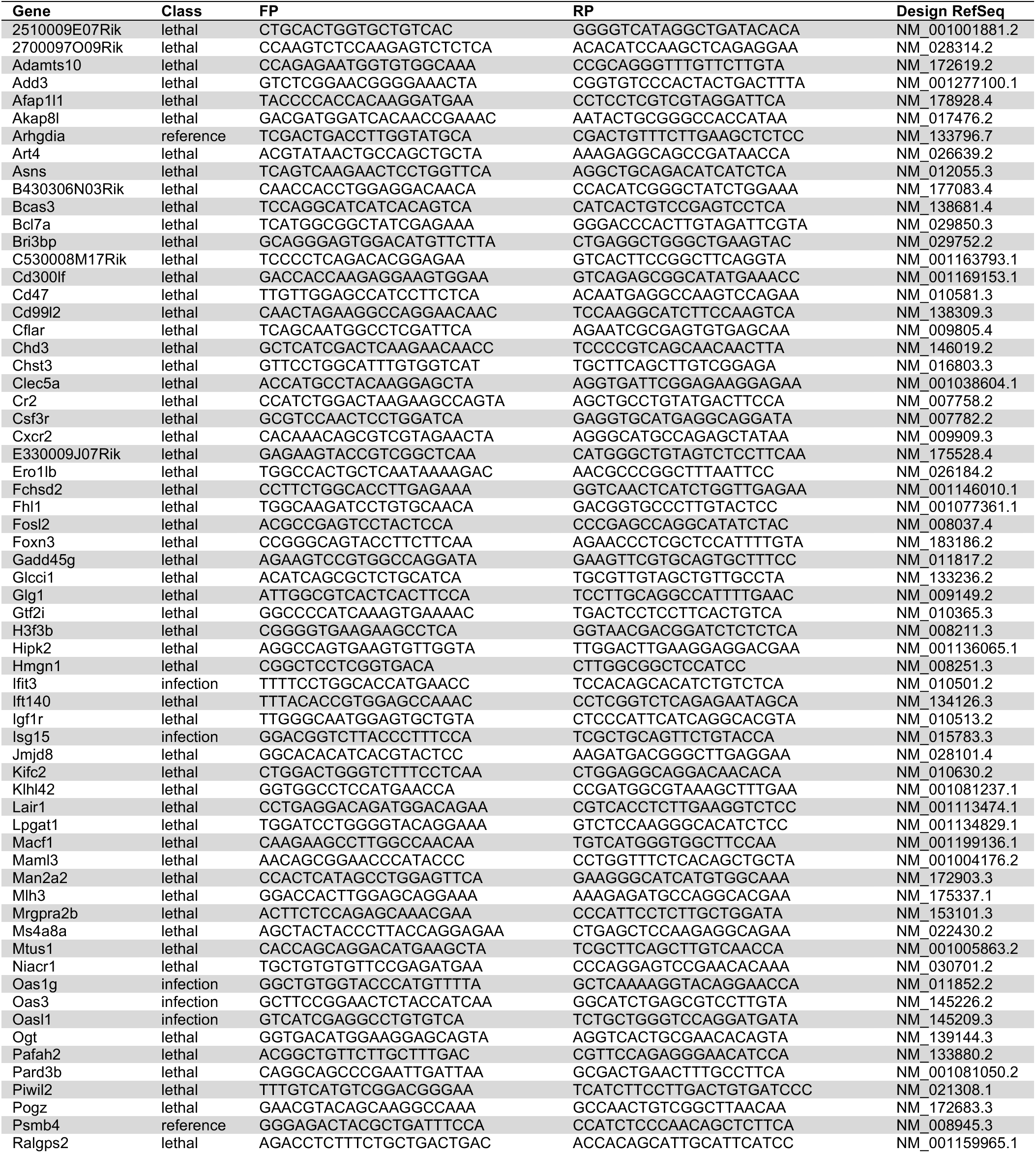
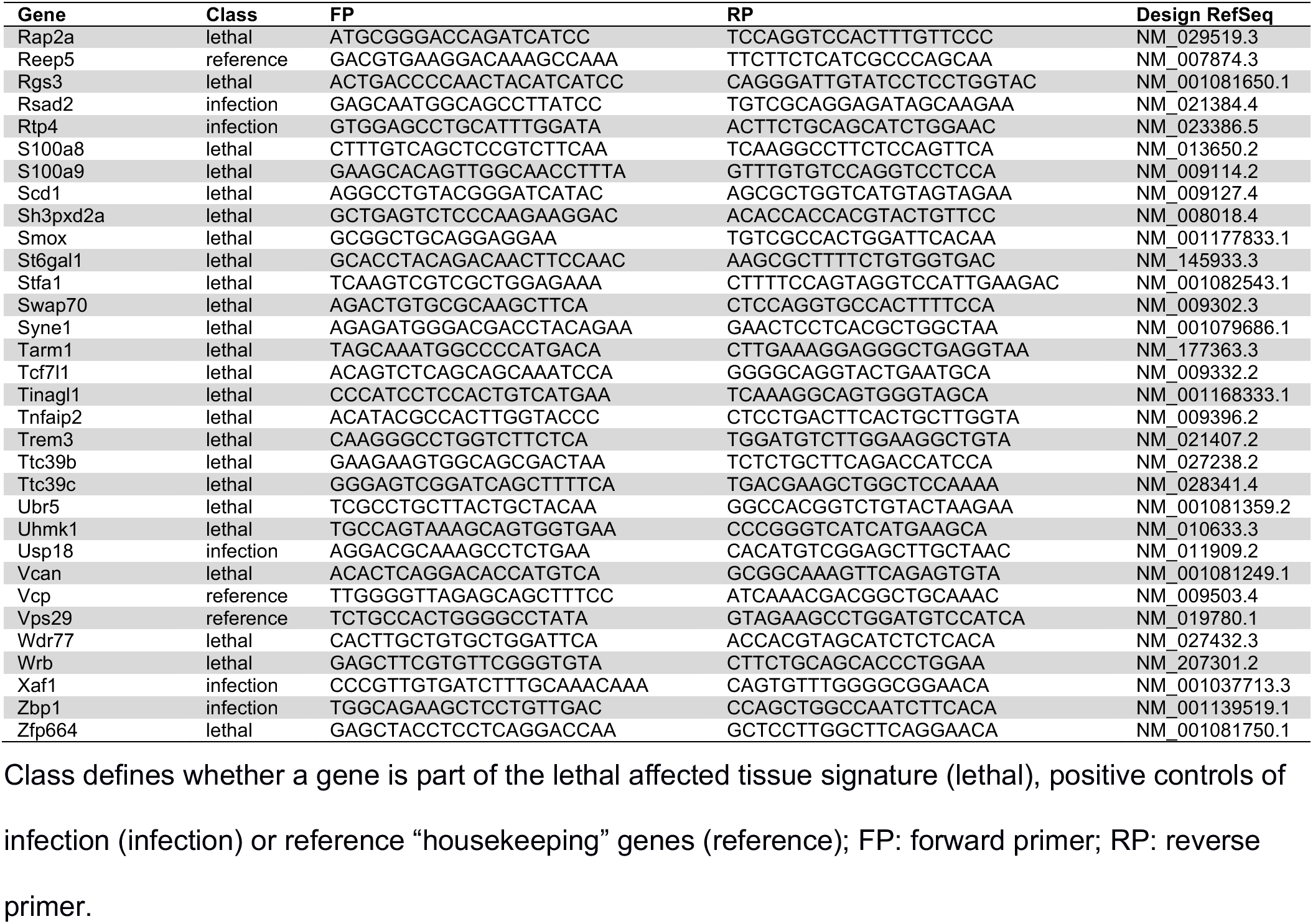
Selected genes whose expression was measured in the blood of mice upon treatment for training and testing model.

